# EzrA coordinates septal organization and nucleoid occlusion in *Staphylococcus aureus*

**DOI:** 10.64898/2026.02.20.707072

**Authors:** Muhammad S. Azam, Tonu Pius, Dominique Missiakas

## Abstract

Envelope biogenesis in *Staphylococcus aureus* is concentrated at the septum and encompasses peptidoglycan synthesis, teichoic acid production, and the targeted secretion and anchoring of YSIRK/GXXS signal peptide-bearing surface proteins. How *S. aureus* spatially restricts these processes to the dividing crosswall remains unclear. EzrA is a membrane-anchored protein structurally related to eukaryotic spectrins that has been implicated in linking cell division to envelope synthesis, yet its precise contribution remains undefined. Here, we show that EzrA spatially couples peptidoglycan synthesis with the surface display of septal proteins. In *ezrA* null cells, newly synthesized peptidoglycan is incorporated in a dispersed pattern rather than confined to the septum, teichoic acid synthesis is increased, and surface protein display is elevated. In wild-type cells, nascent peptidoglycan and nascent Staphylococcal protein A anchoring sites colocalize at the septum; this spatial coupling is lost in the absence of EzrA. *ezrA* null cells also exhibit impaired nucleoid occlusion, with septal ingression over the chromosome leading to chromosome guillotining. Together, these findings suggest that EzrA constitutes a spatial organizing system at the septum, coordinating multiple envelope biogenesis processes while linking septal organization to nucleoid occlusion, a parallel spatial positioning system, to ensure faithful chromosome segregation and cytokinesis.

## Introduction

*Staphylococcus aureus* is an exceptionally adaptable human pathogen whose success depends on a diverse arsenal of virulence factors that mediate host tissue interactions and immune evasion (1, 2). These factors are either secreted as soluble proteins or anchored in the cell envelope; in both cases, their export depends on the Sec secretion pathway (3). *S. aureus* and several other Gram-positive bacteria possess a spatially restricted Sec secretion pathway that targets surface proteins for secretion and anchoring at the dividing septum (4–6). Our laboratory has focused on identifying trans-acting factors that direct proteins containing the YSIRK/GXXS signal peptide to the septum (6, 7). Using a thermosensitive chemical mutagenesis screen, we previously identified PepV as a component of this pathway (8–10). Subsequent investigations revealed that PepV interacts with EzrA and modulates interactions of EzrA with penicillin binding protein 2 (PBP2) and EzrA’s interaction with FtsZ (11), prompting an examination into the contribution of EzrA in coordinating envelope synthesis and cell division.

A distinctive feature of *S. aureus* biology is its cell division program: unlike rod-shaped bacteria that divide along a single axis, *S. aureus* divides in three successive orthogonal planes, cycling through different division axes over consecutive rounds (12). This geometry imposes demands on the spatial coordination of cytokinesis, envelope biogenesis, and chromosome segregation. Bacteria have evolved dedicated spatial positioning systems to meet such demands precisely, ensuring that the divisome assembles at the correct site and that replicated chromosomes are fully segregated before the terminal stage of cytokinesis (13). Yet *S. aureus* lacks two of the most canonical systems, the Min system, which prevents aberrant FtsZ assembly at the cell poles (14, 15), and the ParABS partition system for genome segregation (16–18). ParABS comprises ParA/Soj (a Walker-type ATPase), ParB (a DNA-binding protein), and *parS* (centromere-like sequences near the replication origin) to coordinate genome separation. *S. aureus* lacks a ParA/Soj homolog (19, 20). How, then, does *S. aureus* ensure accurate divisome positioning and faithful chromosome segregation while avoiding bisection of the nucleoid by the closing septum?

The answer appears to lie in a set of organism-specific, partially redundant spatial regulators whose combined action compensates for the absence of these canonical systems. In *S. aureus*, one such system is nucleoid occlusion (NO), mediated by the protein Noc (15). Noc binds conserved chromosomal sequences distributed across the chromosome but excluded from the replication terminus; upon cooperative nucleoprotein complex formation, it blocks FtsZ ring assembly over unsegregated DNA (21, 22). In addition to this role, Noc couples new replication initiation to the completion of chromosome segregation by restricting DnaA activity at origin-proximal Noc-Binding Site (NBS) (21, 22).

Beyond nucleoid occlusion, several additional proteins, including GpsB, DivIVA, and EzrA, ensure spatial regulation at the septum. GpsB stimulates FtsZ GTPase activity and bridges the Z-ring to envelope biogenesis through contacts with EzrA, DivIVA, PBP4, and the wall teichoic acid machinery (23–26). DivIVA is a curvature-sensing protein that localizes to the septum and cell periphery, interacts with multiple divisome components and the chromosome segregation factor SMC, and recruits PcdA to regions of negative membrane curvature that mark the future orthogonal division plane (27–29). Among these regulators, EzrA (Excessive Z rings regulator A) is particularly intriguing. is a membrane-anchored protein proposed to form an arch-like structure above FtsZ filaments (30, 31). It was initially identified in *Bacillus subtilis* as a negative regulator of FtsZ assembly (31, 32). A structural analysis of staphylococcal EzrA revealed striking similarities to eukaryotic spectrins (30). In eukaryotic cells, spectrins function as molecular organizers, coordinating the spatial arrangement of membrane proteins, cytoskeletal elements, and signaling complexes (33, 34). The staphylococcal septum is likewise a site of concentrated biosynthetic activity, serving as the predominant site of peptidoglycan synthesis, teichoic acid (wall and lipoteichoic) biosynthesis, and cell wall protein anchoring (35–37). This report asks whether EzrA plays an analogous organizing role by addressing two questions: whether EzrA help restrict key septal processes, and whether such spatial restriction constitutes a broader positioning system that governs divisome and nucleoid placement, thereby coordinating chromosome segregation with cytokinesis for faithful cell division.

Here, we present evidence that staphylococcal EzrA spatially couples peptidoglycan synthesis with the surface display of septal proteins. Loss of EzrA disrupts this co-localization and is accompanied by increased teichoic acid synthesis, enhanced and dispersed peptidoglycan incorporation, and elevated surface protein display. Cells lacking EzrA fail to maintain proper nucleoid occlusion, allowing septal ingression over the chromosome and resulting in chromosome guillotining. Together, these findings point to a broader role for EzrA in cell division and raise the possibility that EzrA-mediated septal organization and nucleoid occlusion are interdependent, with each process required for the proper spatial positioning of the other.

## Results

### Altered and dispersed synthesis of peptidoglycan in cells lacking EzrA

Unlike rod-shaped bacteria, which synthesize peptidoglycan along both the lateral wall and septum, *S. aureus* concentrates new cell wall incorporation almost exclusively at the division septum (35, 38). If EzrA functions as a septal organizer, its loss should disrupt this spatial restriction, dispersing peptidoglycan synthesis across the cell surface and potentially altering its overall level. To test this, we exploited metabolic labeling with fluorescent D-amino acid analogs, which marks sites of active peptidoglycan incorporation, to assess both the localization and extent of cell wall synthesis in *ezrA* null strain. Wild-type (WT), Δ*ezrA*, and the complemented strain (Δ*ezrA* pCL55-*ezrA*) were pulse-labeled with RADA for 30 minutes, a period corresponding to less than one division cycle, then fixed and stained with BODIPY-FL vancomycin (Fig. 1A). Δ*ezrA* cells showed a marked increase in RADA incorporation relative to WT or complemented controls (Fig. 1B, C), with the RADA signal appearing more broadly distributed across the septal region. As previously reported (39), Δ*ezrA* cells also display pronounced morphological defects, including enlarged cell size and multiple ingressing septa. Two explanations could account for this dispersed labeling pattern: either the peptidoglycan synthesis machinery itself becomes spatially disorganized in the absence of EzrA, consistent with an organizational role for the protein, or the apparent dispersion simply reflects the larger septal area of Δ*ezrA* cells.

**Fig. 1:**
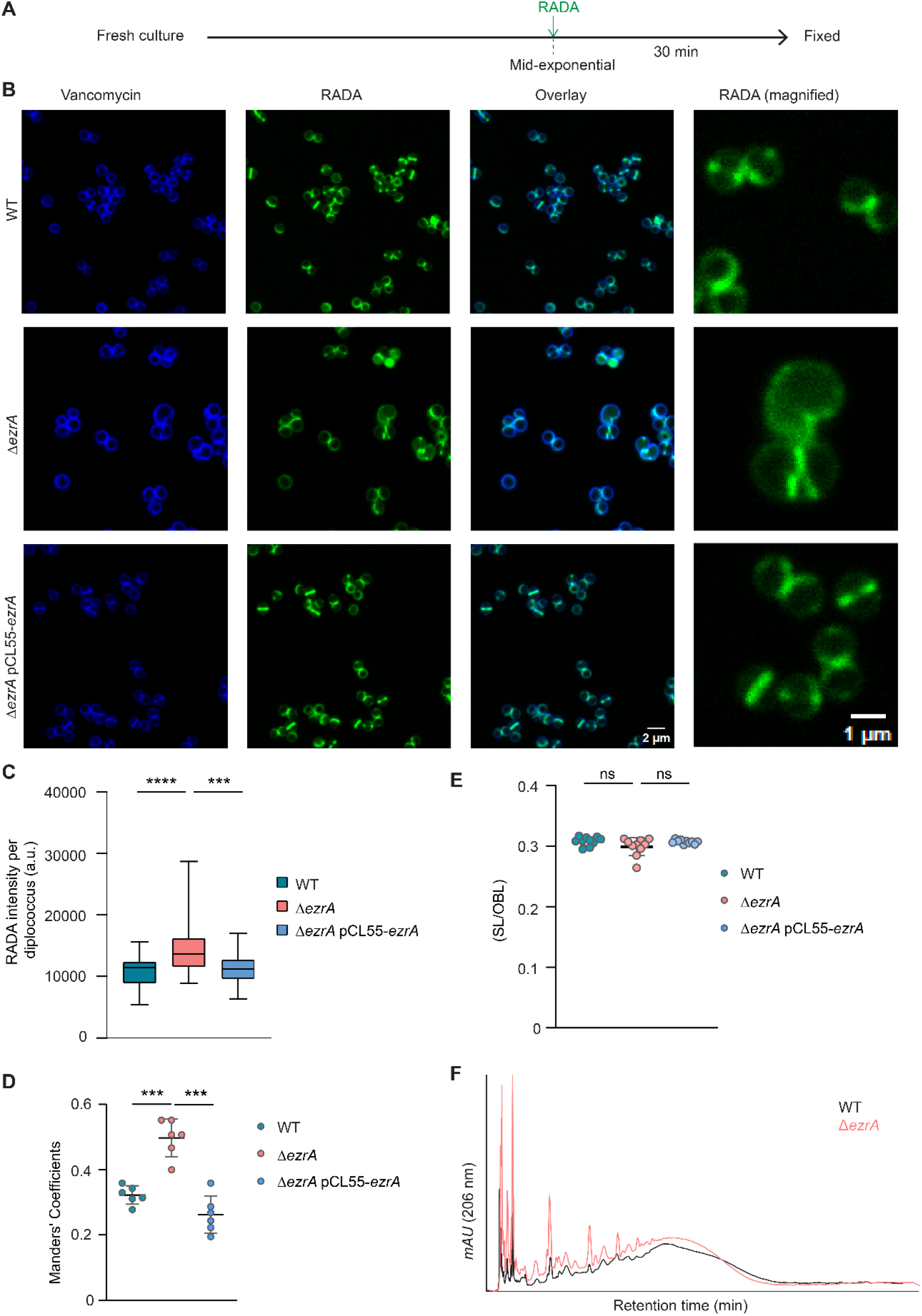
Loss of EzrA leads to enhanced and dispersed peptidoglycan synthesis. (A) Schematic of the labeling experiment. Mid-exponential phase cells were labeled with the fluorescent D-amino acid analog RADA (green) for 30 minutes and fixed. (B) Fluorescence micrographs of fixed cells stained with BODIPY-vancomycin to label the cell wall (blue). (C) Quantification of RADA fluorescence intensity at septal and non-septal (polar) regions in WT (n = 79 cells), Δ*ezrA* (n = 35), and Δ*ezrA* pCL55-*ezrA* (n = 77) cells shown as box-and-whisker plots. Boxes show the interquartile range and median, whiskers show the minimum and maximum values. Data were pooled from two independent experiments. Groups were compared by Welch’s one-way ANOVA. Δ*ezrA* differed significantly from both WT (P < 0.0001) and the complemented strain (P = 0.0003), which did not differ from each other (P = 0.57). *** P < 0.001; **** P < 0.0001. (D) Colocalization of RADA and vancomycin signals, shown as Manders’ coefficients (M values) for the fraction of RADA pixels overlapping with vancomycin pixels. Statistical significance was determined by two-tailed t-test. ***P ≤ 0.001. (E) Ratio of septal length to outer boundary length (SL/OBL) across all three strains assessed from images shown in B. (F) Reversed-phase HPLC of mutanolysin-digested peptidoglycan purified from WT and Δ*ezrA*.

To distinguish between these two possibilities, we first quantified the overlap between RADA incorporation and vancomycin staining (which labels the entire cell periphery) using Manders’ coefficient, the fraction of RADA signal intensity that coincides with vancomycin signal (Fig. 1D). Δ*ezrA* cells exhibited higher coefficients than both WT and complemented strains, indicating that peptidoglycan synthesis is more spatially dispersed in the absence of EzrA. To rule out the possibility that this dispersion is simply due to larger septa, we measured septal lengths and calculated the ratio of septal length to outer boundary length (SL/OBL) across all three strains. The SL/OBL ratio was similar regardless of genotype (Fig. 1E), suggesting that the observed dispersion is not a consequence of septal enlargement. To determine whether increased RADA incorporation in Δ*ezrA* cells (Fig. 1B, C) correlates with increased peptidoglycan content, murein sacculi were purified from WT and mutant bacteria, treated with mutanolysin, and digested preparations separated by reversed-phase high-performance liquid chromatography (HPLC). The resulting elution profiles indicate that Δ*ezrA* samples contain more peptidoglycan than WT (Fig. 1F). Together, these results indicate that loss of EzrA leads to both enhanced and spatially dispersed peptidoglycan synthesis.

### EzrA ensures septal restriction of peptidoglycan synthesis and surface protein display

The staphylococcal envelope displays approximately 24 surface proteins, although the exact number varies by strain (40). One such protein is Staphylococcal protein A (SpA), a YSIRK/GXXS signal peptide–bearing surface protein incorporated into newly synthesized cross-walls by sortase A (41, 42). Since nascent SpA is secreted at the septum and anchored to newly synthesized peptidoglycan, we asked whether SpA anchoring, in extent and in spatial distribution, is altered in the absence of EzrA. Cells were first treated with trypsin to remove surface-exposed proteins, followed by trypsin inhibition (Fig. 2A) (6, 7), and then labeled with fluorescent vancomycin and immunostained for SpA using a monoclonal anti-SpA antibody and a Janelia Fluor 646–conjugated secondary antibody. Δ*ezrA* cells showed markedly more SpA signal than WT, and complementation (Δ*ezrA* pCL55-*ezrA*) restored surface display to WT levels (Fig. 2B, C). Δ*ezrA* cells were also heterogeneous in SpA placement: some diplococci deposited SpA at the septum (Fig. 2B, arrow i), others at ectopic sites (arrow ii), and a third population comprised enlarged cells with no detectable surface SpA (arrow iii).

**Fig 2:**
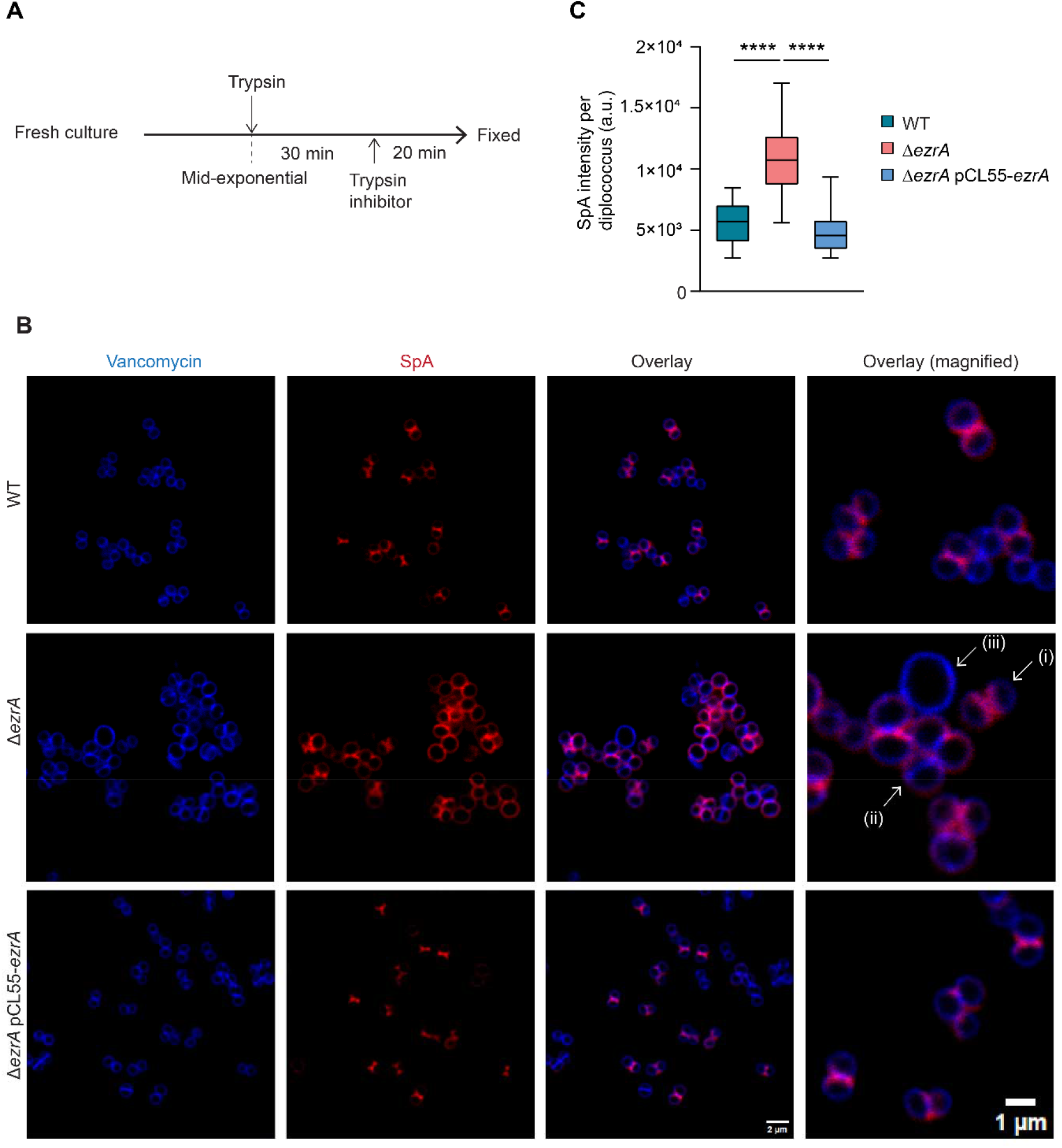
Loss of EzrA increases septal display of newly synthesized surface protein. **A.** (A) Schematic of the labeling experiment. Mid-exponential phase cells were treated with trypsin to remove surface-exposed proteins, then incubated with a trypsin inhibitor to monitor newly synthesized Staphylococcal protein A (SpA). (B) Fluorescence micrographs of fixed cells stained with BODIPY-FL vancomycin (blue) to label the cell wall and immunostained for SpA with a SpA-specific primary antibody and a Janelia Fluor 646–conjugated secondary antibody (red). Arrows (i, ii, iii) in magnified overlay point to distinct cell morphologies. (C) SpA fluorescence intensity per diplococcus, quantified from images in panel A, in WT, Δ*ezrA*, and Δ*ezrA* pCL55-*ezrA* (n = 40 cells per strain) as box-and-whisker plots. Data were pooled from two independent experiments. Groups were compared by Welch’s one-way ANOVA with Games-Howell multiple comparisons test. Δ*ezrA* differed significantly from both WT and the complemented strain (**** P < 0.0001 for each), which did not differ significantly from each other (P = 0.15).

This dispersal of SpA display paralleled the altered peptidoglycan synthesis pattern described above (Fig. 1), raising the possibility that EzrA restricts both processes to the septum, coupling YSIRK/GXXS-dependent protein display to sites of septal peptidoglycan synthesis. To examine this directly, bacteria were first treated with trypsin for 30 min to eliminate surface proteins, next trypsin inhibitor and fluorescent D-amino acid analog RADA were added simultaneously (Fig. 3A). Cells were then imaged for nascent SpA (red) and active sites of peptidoglycan synthesis (green). In WT cells, red and green signals localized predominantly to the septum (Fig. 3B) and Manders’ coefficient analysis confirmed substantial co-localization (Fig. 3C). In Δ*ezrA* cells, both signals were enhanced and dispersed (Fig. 3B), and Manders’ coefficients were significantly reduced relative to WT (Fig. 3C), indicating decreased spatial coupling between surface protein display and peptidoglycan synthesis. Together, these results support a model in which EzrA coordinates septal peptidoglycan synthesis with the localized display of surface proteins during cell division.

**Fig 3:**
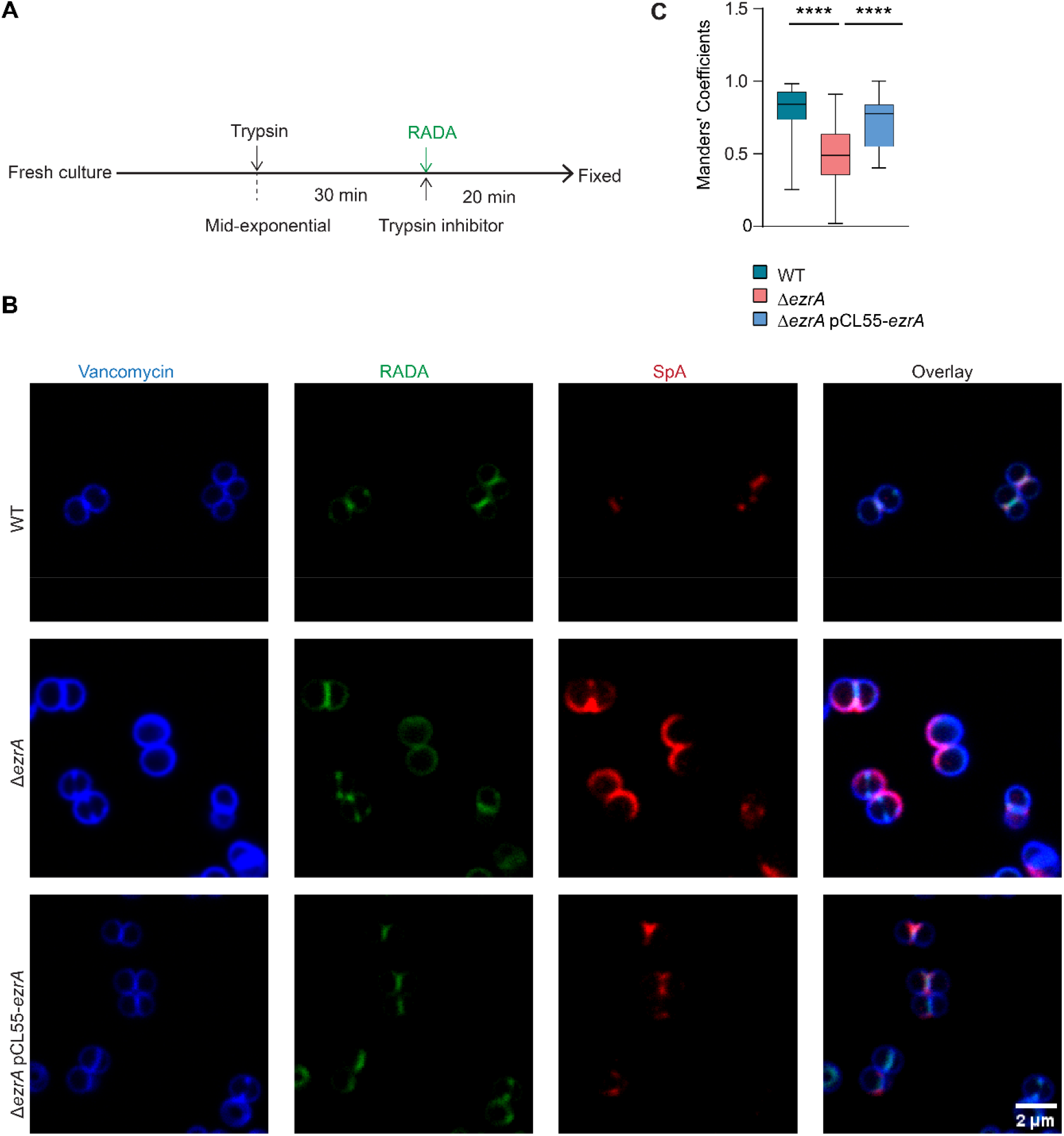
EzrA ensures spatial coupling of peptidoglycan synthesis and protein display at the septum. (A) Mid-exponential phase cells were treated with trypsin to remove surface-exposed proteins, then incubated with a trypsin inhibitor and RADA to monitor newly synthesized SpA and label sites of active peptidoglycan synthesis. (B) Fixed cells were stained with BODIPY-FL vancomycin (blue) and immunostained for SpA (red) as in Fig. 2. (C) Colocalization of SpA (red) and RADA (green) shown as Manders’ coefficients in WT (n = 48 cells), Δ*ezrA* (n = 54), and Δ*ezrA* pCL55-*ezrA* (n = 54) using box-and-whisker plots. Data were pooled from two independent experiments. Groups were compared by ordinary one-way ANOVA, followed by Tukey’s multiple comparisons test. Δ*ezrA* differed significantly from both WT and the complemented strain (**** P < 0.0001 for each). The complemented strain differed slightly from WT (P = 0.029, not marked on the plot).

EzrA-dependent septal organization may also shape the final step of cytokinesis: the physical separation of daughter cells. *S. aureus* daughter cells separate through a rapid “popping” event in which the peripheral ring joining them cracks open asymmetrically, leaving the two cells transiently tethered by a hinge (43). This crack-propagation mode of separation is uncommon among bacteria. It has been described only in staphylococci, their coccoid relatives in the genus *Macrococcus*, and members of the Actinobacteria, and its determinants remain poorly understood (44). Because cracking occurs on a millisecond timescale, scanning electron microscopy (SEM) has captured only pre- and post-cracking states (43). A previous report showed that, in the pre-cracking state, small perforations appear in the peripheral ring, likely resulting from mechanical stress combined with hydrolase activity (43). These perforations serve as initial failure points from which the crack propagates around the circumference, leaving a hinge roughly opposite the initiation site (43). Consistent with these reports, we did not detect intermediate states in WT cells using SEM (Fig. 4A). In Δ*ezrA* cells, however, we observed cells in which the crack appeared to have paused or slowed around the circumference (Fig. 4A).

**Fig 4:**
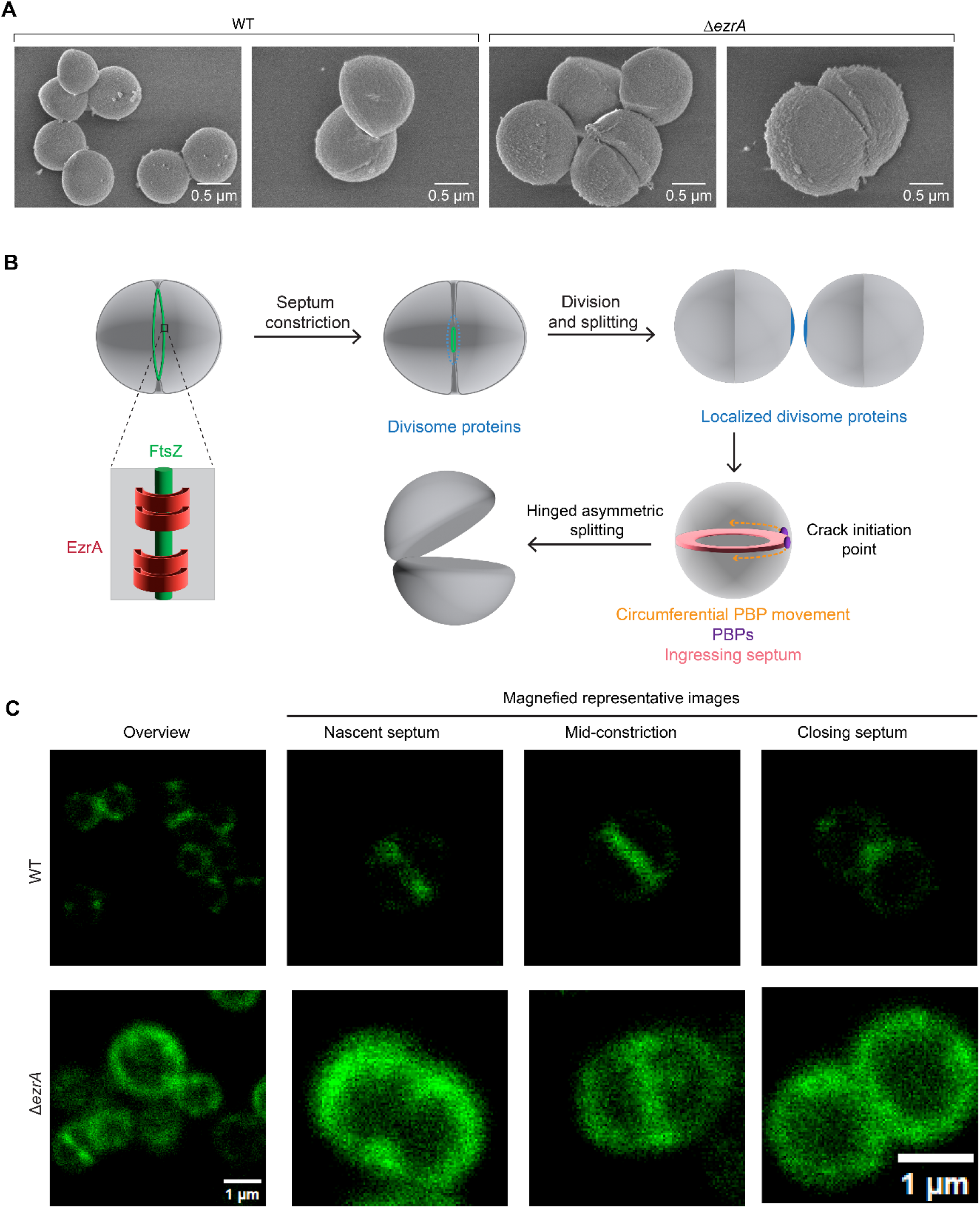
Concentration of septal components may explain the initiation of ‘crack propagation’ during popping that drives staphylococcal cell separation. **(A)** Scanning electron microscopy (SEM) of WT and Δ*ezrA* cells (B) EzrA ensures concentration of PBPs at the ingressing septum during cytokinesis. During late cytokinesis, divisome components remain concentrated at a discrete site within the newly synthesized envelope. This localized concentration may serve as a point for initiation of the next round of septum formation. (C) Representative STED micrographs of WT and Δ*ezrA* cells labeled with Bocillin-FL to visualize penicillin-binding proteins (PBPs). In WT cells, PBPs concentrate at a single focus during late cytokinesis, consistent with the model in panel B.

The site at which perforations first form, and thus where the crack initiates, has been presumed to be random (43). The molecular organization model presented here suggests an alternative. If the septal machinery were progressively concentrated as division proceeds, it would define the site where septal PBPs begin circumferential synthesis, making that point the oldest region of the septum and the hinge of the youngest (Fig. 4B). Hydrolases resolving the septum might then act first at this same site, positioning it as the origin of the crack. We therefore propose that EzrA-mediated organization focuses membrane proteins at the septum through early and mid-constriction, culminating in their accumulation at a discrete point on the closing septum, a site that could both initiate crack propagation and seed the next round of division (Fig. 4B). To test this, mid-exponential cells were pulse-labeled with the fluorescent penicillin derivative Bocillin-FL, which forms a stable acyl-enzyme intermediate with PBPs, and imaged by stimulated emission depletion (STED) microscopy. In WT cells, PBPs concentrated at the ingressing septum and, in late cytokinesis, resolved to a distinct focus at the closing septum, consistent with the model (Fig. 4C). In Δ*ezrA* cells, this distribution was disrupted: PBPs were neither confined to the dividing septum nor focused into a discrete point at late cytokinesis (Fig. 4C). These data suggest that EzrA concentrates PBPs into a discrete focus at the closing septum, positioning the septal machinery to specify where the peripheral ring first fails. Whether the crack initiates at this focus and whether hydrolase activity is recruited there, remains to be shown. Resolving this will require correlating PBP focus position with crack initiation site in individual dividing cells, which we will pursue in future work.

### Enhanced teichoic acid synthesis in the absence of EzrA

The data presented above support a role for EzrA in organizing both cytoplasmic and cell envelope-associated components of the divisome. Because wall teichoic acid (WTA) and lipoteichoic acid (LTA) are likewise assembled septally and mature outward, we asked whether their abundance is altered in the absence of *ezrA* (45, 46). Murein sacculi isolated from WT, Δ*ezrA*, and complemented strains were alkali-treated to release WTA polymers that were resolved by polyacrylamide gel electrophoresis (PAGE) and visualized by alcian blue-silver staining (Fig. 5A). The Δ*ezrA* strain displayed increased WTA abundance compared to both controls (Fig. 5A), a finding consistent with fluorescence microscopy using wheat germ agglutinin (WGA) conjugated to Alexa Fluor 488 (Fig. 5B, C); WGA is a lectin that recognizes GlcNAc-containing surface structures, including peptidoglycan and glycosylated WTA and LTA (47). Immunoblot analysis similarly revealed increased LTA content in the Δ*ezrA* strain compared to controls (Fig. 5D). Together, these results indicate that both WTA and LTA are overproduced in the absence of EzrA. Whether this reflects increased biosynthetic enzyme abundance or altered localization of existing enzyme pools remains to be determined.

**Fig 5:**
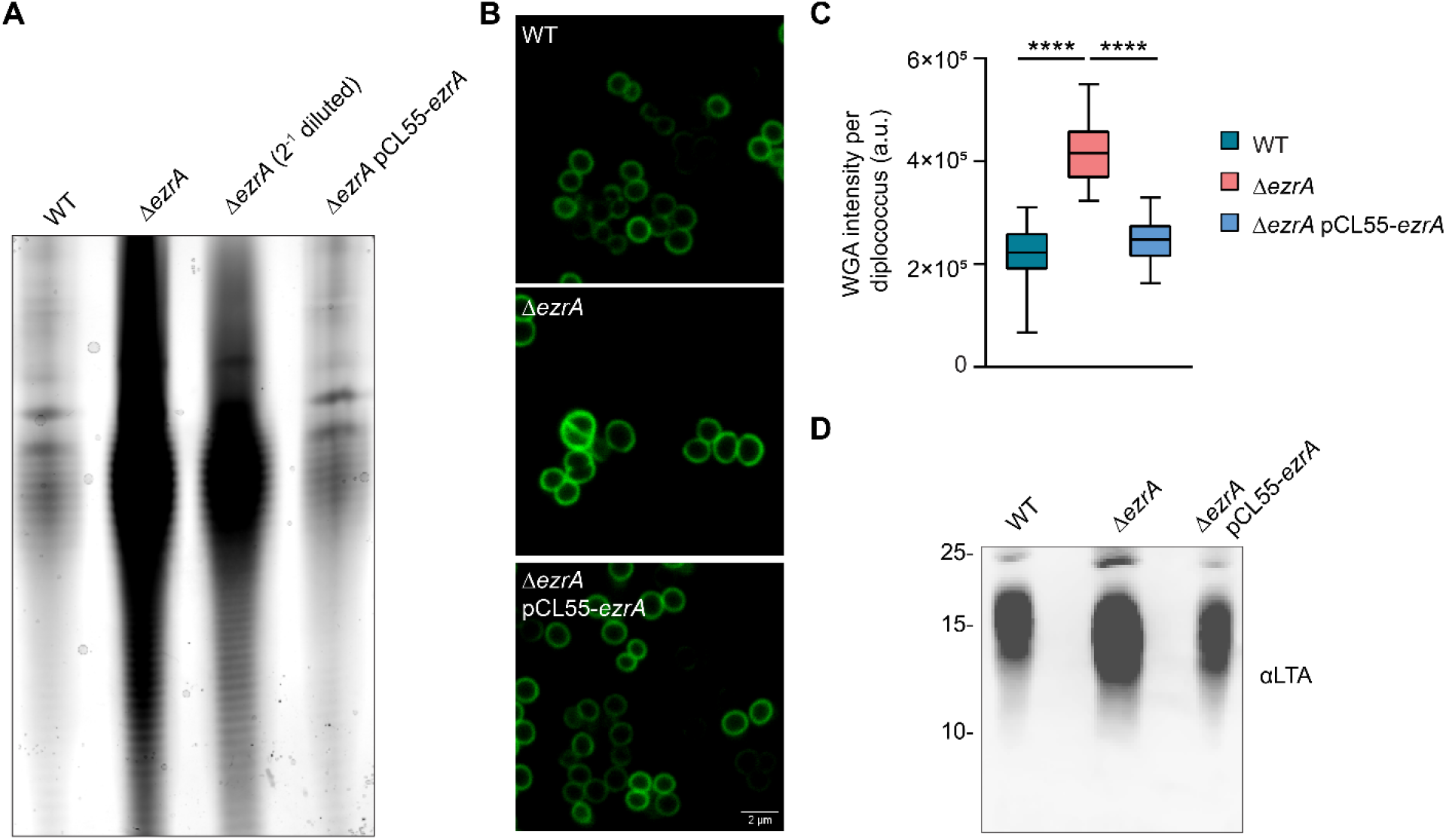
Loss of EzrA increases teichoic acid synthesis. (A) WTA extracted from WT, Δ*ezrA*, and Δ*ezrA* pCL55-*ezrA*, resolved by PAGE, and visualized by alcian blue-silver staining. (B) Fluorescence micrographs of WT, Δ*ezrA*, and ΔezrA pCL55-*ezrA* cells stained with fluorescent WGA. (C) WGA fluorescence intensity per diplococcus in WT, Δ*ezrA*, and Δ*ezrA* pCL55-*ezrA* (n = 40 cells per strain) shown as box-and-whisker plots. Data were pooled from two independent experiments. Groups were compared by Welch’s one-way ANOVA with Games-Howell multiple comparisons test. Δ*ezrA* differed significantly from both WT and Δ*ezrA* pCL55-*ezrA* (**** P < 0.0001 for each). The complemented strain was slightly higher than WT (P = 0.044). * P < 0.05; **** P < 0.0001. (D) LTA levels in WT, Δ*ezrA*, and Δ*ezrA* pCL55-*ezrA*, assessed by immunoblotting with anti-LTA (α-LTA) antibodies.

### Nucleoid occlusion is affected in the absence of EzrA

Across the bacterial kingdom, cytokinesis, envelope biogenesis, and chromosome segregation are executed as a single coordinated program rather than as independent processes (48–51). This coordination depends on membrane-associated proteins that bridge cytoplasmic DNA to the cell membrane. Nucleoid occlusion illustrates this principle (49, 52, 53). Noc, a ParB-family protein that binds both chromosomal DNA and the membrane, blocks assembly of the division machinery over the nucleoid and thereby prevents chromosome bisection by the closing septum (15, 21, 54). Unlike SlmA in *E. coli*, Noc does not appear to act through direct contact with FtsZ; instead, it inhibits division by tethering NBS-bound DNA to the membrane via an N-terminal amphipathic helix (22). In *S. aureus,* loss of Noc impairs orthogonal FtsZ ring formation, leading to Z-ring assembly over the nucleoid and chromosome guillotining (15).

Because loss of EzrA also results in impaired orthogonal FtsZ ring formation, we asked whether EzrA is required for proper nucleoid occlusion. To address this, we constructed a *noc*′*–sfgfp* fusion under an anhydrotetracycline inducible promoter (P*_itet_*) and integrated it at the *geh* locus using the pCL55 vector system that utilizes the site-specific recombination system of staphylococcal phage L54a (55). Mid-exponential phase cells were pulse induced with anhydrotetracycline to transiently express Noc-sfGFP fusion protein (Fig. 6A). In WT cells, Noc was distributed throughout the cell; in Δ*ezrA* cells, the green signal was more intense and in a subset of cells appeared enriched near the cell periphery. Here, peptidoglycan was labeled with Cy5-DA, a fluorescent D-amino acid probe (Fig 6A). Immunoblot analysis of mid-exponentially grown WT, Δ*ezrA*, and complemented cells confirmed that Noc abundance is increased in the Δ*ezrA* strain compared to both controls (Fig. 6B, C), suggesting that nucleoid occlusion is not operating normally in the absence of EzrA.

**Fig. 6.**
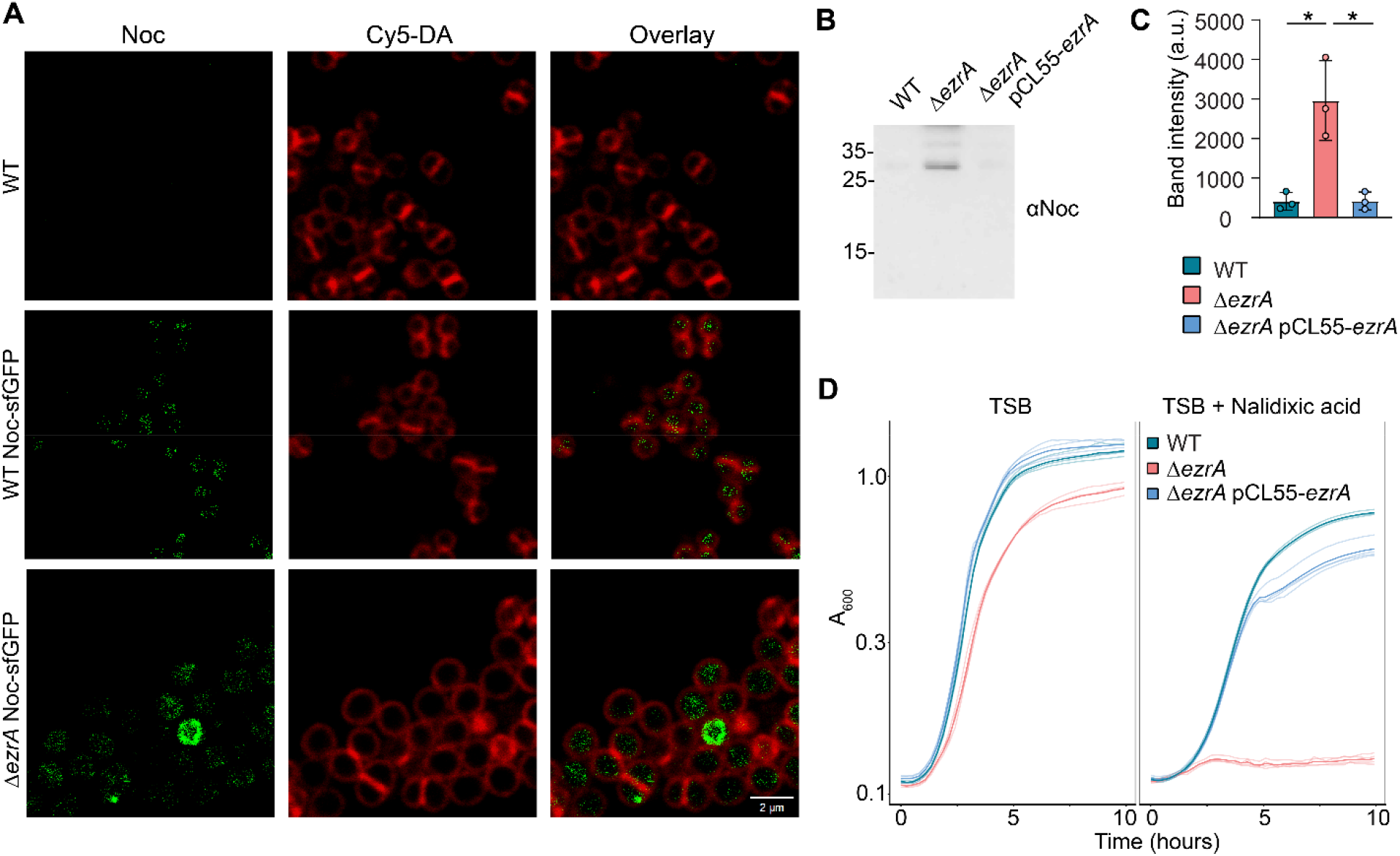
Nucleoid occlusion is altered in the absence of EzrA. (A) Representative fluorescence micrographs of WT cells (control) and of WT and Δ*ezrA* cells expressing itet-inducible *noc*–sfGFP from mid-exponential phase cultures induced with anhydrotetracycline for 1 h. (B) Immunoblot of Noc in WT, Δ*ezrA*, and complemented ΔezrA pCL55-*ezrA* strains. (C) Quantification of Noc immunoblot signal from three independent experiments. Statistical significance was assessed using a two-tailed *t*-test (*, *P* ≤ 0.05). (D) Growth curves of WT, Δ*ezrA*, and complemented ΔezrA pCL55-*ezrA* strains in TSB containing nalidixic acid at 37°C with shaking. Overnight cultures were diluted into the medium at time zero, and absorbance at 600 nm (A_600_) was measured at the indicated time points.

If nucleoid occlusion is impaired, the consequences should be most apparent when chromosome replication and segregation are transiently uncoupled from division, when Noc function is most critical (49). We therefore challenged cells with a sublethal concentration of nalidixic acid, a quinolone that targets DNA gyrase (56) and slows replication elongation (57), placing the occlusion machinery under demand. Δ*ezrA* cultures were markedly more sensitive than WT under these conditions, and growth was largely restored by complementation (Fig. 6D).

Defective nucleoid occlusion raises the possibility that septal constriction occurs over the chromosome, leading to DNA damage, inefficient separation, and the formation of anucleated or partially nucleated cells. To test this possibility, mid-exponential phase WT, Δ*ezrA*, and complemented cells were fixed and stained with Hoechst to visualize DNA and with BODIPY FL conjugated vancomycin to label peptidoglycan (Fig. 7). Δ*ezrA* cells included enlarged cells that were either partially or completely devoid of nucleoid material (Fig 7, red and yellow arrows, respectively in the overlay image of Δ*ezrA* cells). Approximately 2.3% of Δ*ezrA* cells lacked a nucleoid entirely, compared with ∼0.3% of WT and ∼0.04% complemented cells. This increase is consistent with previous reports of DNA guillotining and suggests that EzrA helps prevent Z-ring assembly over unsegregated or segregating chromosomes (15, 20). To investigate whether aberrant Z-ring assembly in Δ*ezrA* cells contributes to these phenotypes, FtsZ-eYFP was ectopically expressed from the P*spac* promoter in mid-exponential-phase WT and Δ*ezrA* cells (Fig. 8). In WT cells, FtsZ-eYFP rings assembled predominantly at midcell, flanked by DNA on both sides. In contrast, Δ*ezrA* cells displayed several aberrant patterns (Fig. 8): some contained more than one ingressing FtsZ ring (red arrow), others formed rings at off-center positions (green arrow), and still others assembled rings near midcell but produced a compartment with less nucleoid DNA (orange arrow). These phenotypes are consistent with chromosome bisection by the division septum, previously described as DNA guillotining (15, 49, 58).

**Fig. 7.**
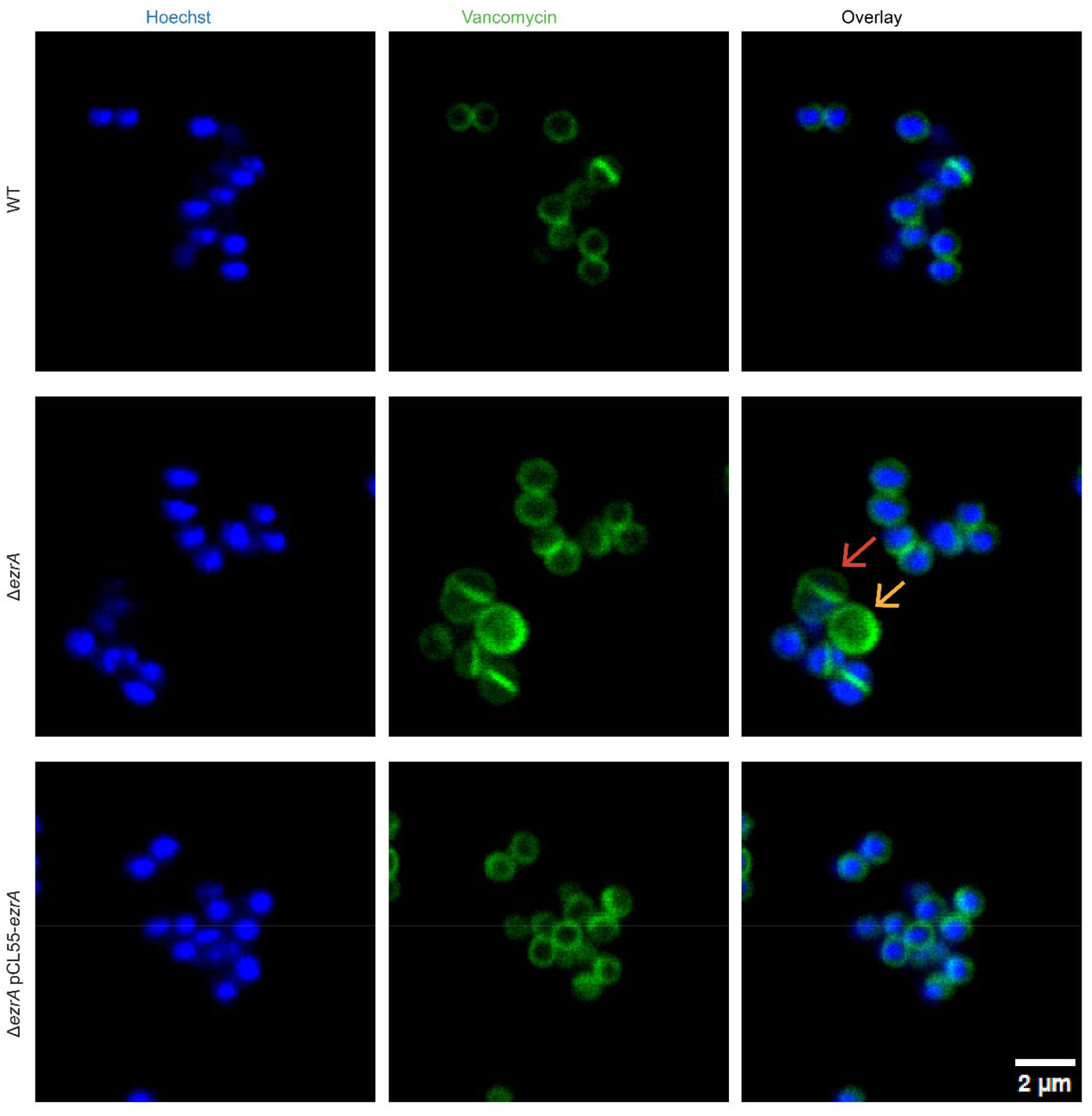
Loss of EzrA promotes the formation of anucleate and partially nucleated cells. Representative fluorescence micrographs of fixed mid-exponential phase WT, Δ*ezrA*, and Δ*ezrA* pCL55-*ezrA* cells stained with fluorescent vancomycin (cell wall) and Hoechst (DNA). In the Δ*ezrA* overlay, red and yellow arrows indicate partially nucleated and anucleate cells, respectively.

**Fig 8.**
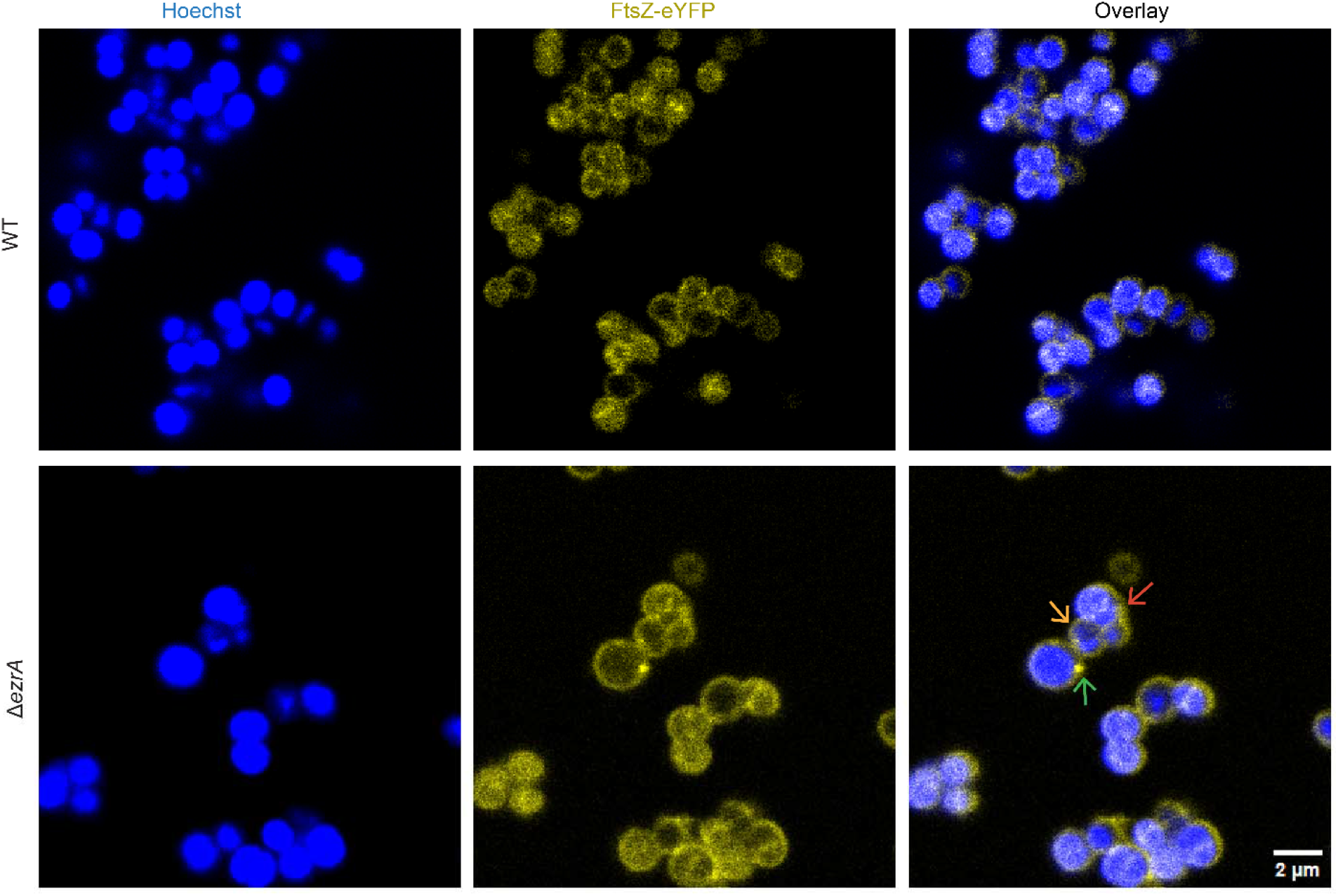
Aberrant FtsZ ring formation in the absence of EzrA. Localization of FtsZ-eYFP in WT and Δ*ezrA* cells. eYFP-tagged FtsZ was induced with IPTG in mid-exponential phase cells, which were then fixed and stained with Hoechst (DNA). Red, green and orange arrows point to Δ*ezrA* cells with aberrantly placed FtsZ rings.

To assess DNA breakage, we performed a terminal deoxynucleotidyl transferase dUTP nick-end labeling (TUNEL) assay. In this assay, permeabilized cells are incubated with terminal deoxynucleotidyl transferase and fluorescently labeled dUTP. Δ*ezrA* cells exhibited robust TUNEL signal, whereas WT cells showed no detectable labeling, indicating the presence of DNA strand breaks specifically in the absence of EzrA (Fig. 9). Taken together, these phenotypes closely resemble those of *noc* null mutants and indicate that EzrA is required for proper nucleoid occlusion. In the absence of EzrA, Noc fails to effectively exclude the division machinery from the nucleoid, resulting in chromosome guillotining, DNA strand breaks, loss of genomic integrity, and reduced cell viability.

**Fig. 9.**
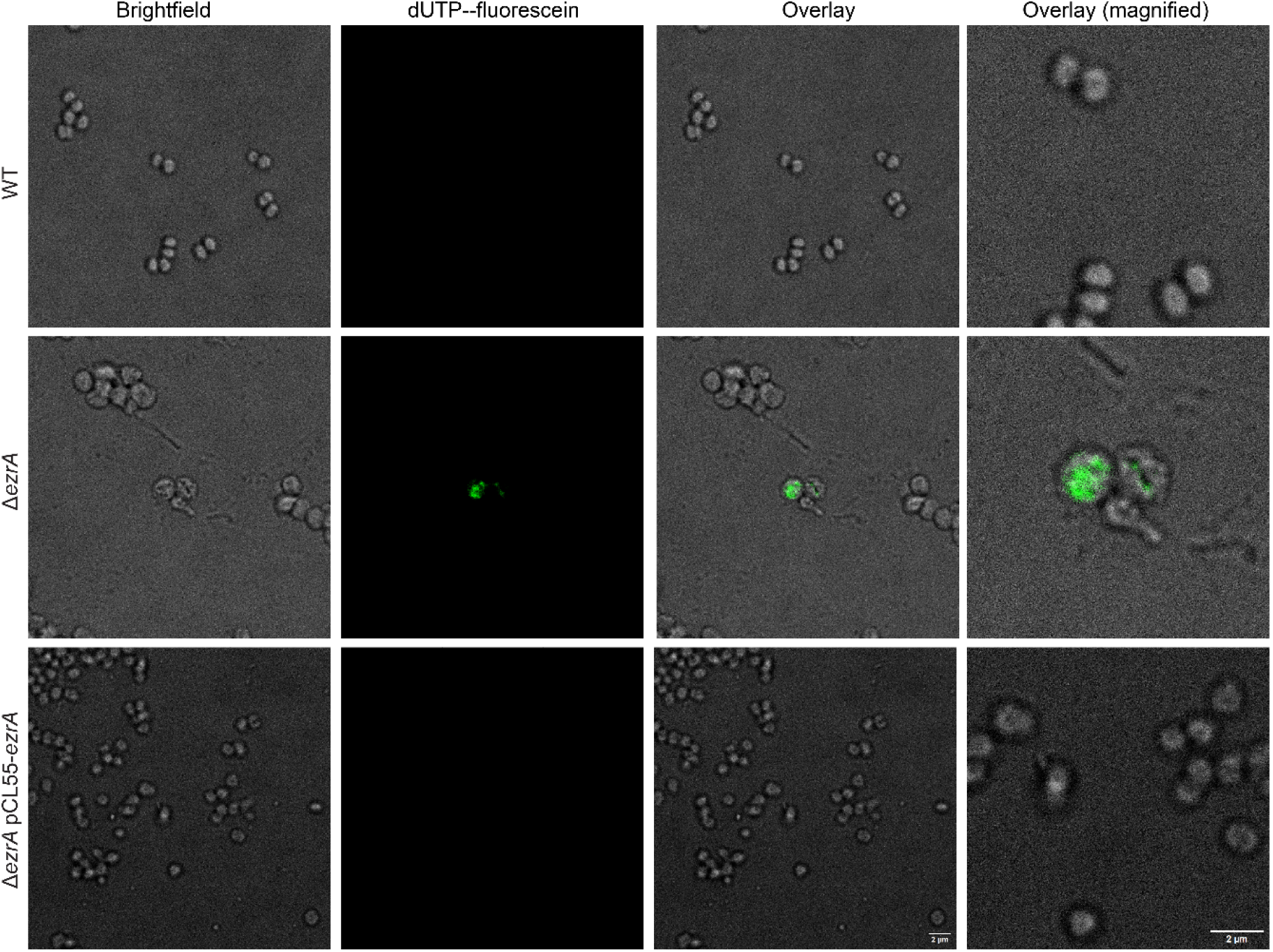
Deletion of *ezrA* leads to DNA fragmentation. DNA breaks were detected using a TUNEL assay in which membrane-permeabilized cells were incubated with terminal deoxynucleotidyl transferase and fluorescein-labeled dUTP. Representative images are shown for WT, Δ*ezrA* and Δ*ezrA* pCL55-*ezrA* cells. Panels show brightfield, fluorescein-dUTP labeling of DNA ends, and merged (same scale and magnified) images, respectively.

## Discussion

Binary fission requires a cell to solve two spatial problems at once: the divisome must assemble at a single site and be confined there, and the replicated chromosome must be cleared from that site before constriction is complete. In rod-shaped model organisms, both challenges are addressed largely through negative regulation: the Min system restricts Z-ring assembly to midcell by maintaining a polar gradient of a septation inhibitor (59, 60) while nucleoid occlusion prevents septum formation over unsegregated DNA (49, 53). Clearance of the chromosome from the furrow is achieved partially by the ParABS partitioning system, in which ParB spreads from centromere-like *parS* sites to form a partition complex that translocates across the nucleoid by stimulating ATP hydrolysis in a ParA gradient, depleting ParA behind it and biasing its own displacement towards the pole (16, 61, 62). *S. aureus* has taken a different approach. It lacks the Min system entirely (14), and the ParABS chromosome partitioning system is incomplete: ParB (Spo0J) and SMC are present, but no ParA/Soj homologue is encoded (14, 20, 63, 64).

Against this background, we identify EzrA as a factor in cell division and chromosome separation. We propose that EzrA functions not only as a scaffold that organizes the placement of division components and the enzymes responsible for septal envelope synthesis (30, 39, 65), but also as a factor required for effective establishment of nucleoid occlusion. The data support a model in which these two systems, nucleoid occlusion and septal organization, differ in the kind of control they exert. Nucleoid occlusion, like the well-studied Min system, is negative and permissive: it does not specify the division site but leaves midcell as the only position at which functional FtsZ assemblies are allowed to form. The role we propose for EzrA is non-negative. In its absence, FtsZ rings form aberrantly, septal processes lose both their spatial restriction and their co-localization with one another, and occlusion is no longer effectively imposed. Together, these observations suggest that two interdependent systems orchestrate staphylococcal division: negative nucleoid occlusion and non-negative septal organization (Fig. 10).

**Fig. 10.**
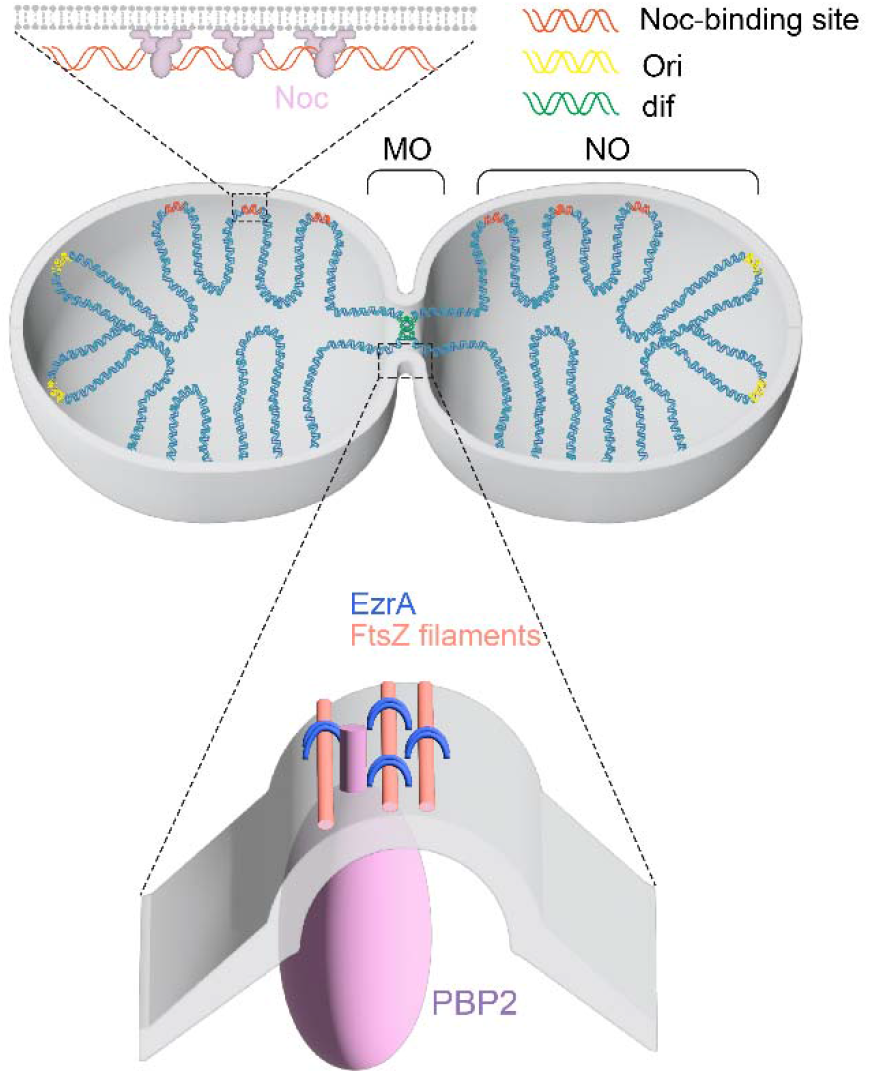
Proposed model for the dual role of EzrA in septal organization and nucleoid occlusion in *S. aureus*. A dividing *S. aureus* cell is shown with its replicated chromosomes (blue). The origin (Ori; yellow) and the *dif* site (green) are indicated; the *dif*-proximal terminus region remains within the furrow as the septum closes. Noc (purple) binds Noc-binding sites (orange) distributed across the chromosome and tethers the DNA to the membrane (upper inset), preventing septum formation over the nucleoid (NO, nucleoid occlusion zone; MO, molecular organization). At the septum (lower inset), EzrA (blue) associates with FtsZ filaments (pink) and the peptidoglycan synthase PBP2 (purple). We propose that EzrA is required both to organize septal proteins and to establish effective nucleoid occlusion. When these processes are coordinated, septal envelope synthesis proceeds correctly, the Z-ring is placed in the correct plane, and replicated chromosomes are fully segregated before septum closure, protecting them from guillotining.

A question left open by this work, and by the field more broadly, concerns the mechanistic basis of guillotining. Is DNA breakage caused directly by the ingressing septum, or can replication perturbation contribute independently? Attributing the Δ*noc* phenotype, or the Δ*ezrA* phenotype reported here, exclusively to mispositioned cytokinesis is likely an oversimplification, since Noc also restrains replication initiation: Δ*noc* cells over-initiate replication, *dnaA* mutations that suppress over-initiation partially suppress the division defects, and DnaA overexpression exacerbates both (21). The most direct causal evidence that the septum itself is the cutting agent comes from *E. coli*, where SOS induction in dimer-resolution mutants follows an apparently normal division event, is abolished when division is blocked, and is accompanied by localized degradation of the terminus region around *dif* (66). Distinguishing mechanical cleavage from replication-derived breakage in *S. aureus* will ultimately require the equivalent experiment: testing whether the DNA damage in Δ*ezrA* cells persists when septum closure is prevented.

A second question concerns penetrance. Our data indicate that EzrA imposes spatial order on septation, yet only a subpopulation of Δ*ezrA* cells displays any given phenotype. Partial penetrance is in fact characteristic of this system, indeed many Δ*noc* cells also divide normally, which has been taken as evidence for additional Z-ring placement factors (15, 21) and the severe defects of a Δ*pcdA* Δ*noc* double mutant indicate that PcdA/DivIVA and Noc operate in parallel (29). Partial penetrance is therefore the expected output of redundant placement systems, in which loss of one factor is buffered in most cells and becomes penetrant only when a second fails within the same division cycle. We favor an additional, non-exclusive hypothesis specific to EzrA: that its septal function and nucleoid occlusion are interdependent rather than sequential. In the absence of EzrA, nucleoid occlusion is not effectively established, permitting guillotining and DNA loss; the severity of the resulting phenotype would then depend on the amount of DNA lost and the identity of the lost segment. Resolving this will require moving beyond population-level snapshots to single-cell lineage tracking, which would reveal whether affected cells are those that inherited a compromised chromosome from the preceding division, and establish whether guillotining in a coccus is a self-propagating failure rather than an isolated event.

## Materials and Methods

### Media, growth conditions, bacterial strains, and plasmids

*S. aureus* strain RN4220 was used as the wild-type (WT) strain to generate isogenic derivatives used for all experiments (67). Strains were grown in tryptic soy broth (TSB) or on tryptic soy agar (TSA) plates and, when appropriate, supplemented with erythromycin or chloramphenicol at a final concentration of 10 μg/mL. *Escherichia coli* XL10 cells were used for cloning and were cultured in lysogeny broth (LB) or on LB agar supplemented with 100 μg/mL ampicillin for plasmid selection. Deletion of *ezrA* gene was achieved through allelic replacement by pKOR1, an *E. coli-S. aureus* shuttle vector as previously described (7, 68) to yield strain RN4220 Δ*ezrA* (SA423). The plasmid pCL55-itet-*noc*-sfGFP was constructed by amplifying the pCL55-itet-sfGFP backbone using primers OSA1680/OSA1681, and the *noc* gene was amplified from RN4220 genomic DNA using primers OSA1686/OSA1687. The resulting fragments were assembled using NEB HiFi DNA Assembly Master Mix (69). The pCL55-itet-*noc*-sfGFP plasmid was introduced by electroporation into RN4220 and RN4220 Δ*ezrA* strains to integrate the itet-*noc*-sfGFP construct at the *geh* locus (55) yielding strains SA583 and SA584, respectively. Expression of Noc-sfGFP was induced by addition of anhydrotetracycline to a final concentration of 100 ng/mL for 1 hour in mid-exponential-phase cultures. Expression of FtsZ-eYFP was ectopically induced in mid-exponential cultures by addition of IPTG to a final concentration of 50 µM using plasmid pCQ11-FtsZ-eYFP (70), obtained from Dr. Foster, University of Sheffield. For growth curve analysis, nalidixic acid was added to liquid cultures at a final concentration of 30 µg/mL.

### Fluorescence microscopy

For fluorescence microscopy, bacterial cells were fixed and immunostained largely as described (7, 71). Briefly, for analysis of surface protein display (SpA immunostaining), cells were harvested, washed with phosphate-buffered saline (PBS), and treated with trypsin (0.5 mg/mL) for 1 h at 37°C. Cells were then washed twice with PBS and resuspended in 900 μL fresh tryptic soy broth (TSB) supplemented with soybean trypsin inhibitor (1 mg/mL), followed by incubation at 37°C for the indicated times. Where indicated, RADA and Cy5-DA were added during this incubation at a final concentration of 100 nM. Cells were fixed for 20 min in 2.5% paraformaldehyde and 0.006% glutaraldehyde, washed twice with PBS, and immobilized on poly-L-lysine–coated glass slides. Immobilized cells were blocked with 3% (wt/vol) bovine serum albumin in PBS for 30 min at room temperature and incubated with SpA-specific primary antibody. After washing with PBS, samples were incubated with JF646-conjugated IgG secondary antibody for 3 h at room temperature. Where indicated, slides were washed with PBS and incubated with BODIPY-FL vancomycin (1 µg/mL, Invitrogen), Alexa Fluor-conjugated wheat germ agglutinin (1 µg/mL), and Hoechst (50 µg/mL) for 10 min. Slides were mounted using ProLong Diamond Antifade (Invitrogen) to minimize photobleaching. Images were acquired using a Leica Stellaris confocal microscope. All microscopy experiments were performed at least twice. 2D STED imaging was conducted on the same microscope, with a 592 nm STED depletion laser applied in addition to the excitation laser.

To quantify SpA signal intensity, measurements were performed manually using ImageJ, with intensity calculated per diplococcus as described previously (72). Diplococci were defined as pairs of daughter cells exhibiting a near-complete or fully formed septum. Colocalization between fluorescence channels was quantified using Manders’ overlap coefficients, which estimate the fraction of signal in one channel that overlaps with signal in the other based on thresholded pixel intensities (73). Two approaches were used. For whole-field analysis (Fig. 1C), coefficients were calculated from entire images containing all bacterial cells in the field using the JACoP plugin for ImageJ (72, 74). For single-cell analysis (Fig. 3), coefficients were calculated with a custom Python script. Briefly, individual cells were segmented from the fluorescent vancomycin channel, which was used to define cell boundaries, and Manders’ coefficients were calculated for each cell from raw pixel intensities within the corresponding mask (73). Thresholds were determined using the Costes method (75). Only cells with above-threshold signal in both RADA and SpA channels were included. This analysis was performed using scikit-image 0.26 (76).

The ratio of septal length to outer boundary length (SL/OBL) was calculated for randomly selected diplococci that had completed septal ingression but had not yet separated. Binary images were analyzed using a custom Python script to identify the largest contour. The outer boundary length (OBL) was calculated as the perimeter of the contour, and septal length (SL) was estimated as the minor axis length. Perimeters were calculated using Ramanujan’s first approximation for ellipse circumference, 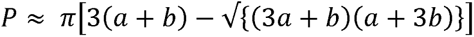 where *a* and *b* denote the semi-major and semi-minor axes, respectively (77).

Statistical analysis was performed in R (78). Normality was assessed using the Shapiro-Wilk test, and homogeneity of variance using the Brown-Forsythe and Bartlett tests (79, 80). Groups were compared by one-way ANOVA followed by Tukey’s multiple comparisons test or, when variances were unequal, by Welch’s ANOVA followed by the Games-Howell multiple comparisons test (P < 0.05 was considered statistically significant) (81–83). Sample sizes and number of biological replicates are indicated in the figure legends. All R and Python scripts used in this study are available on GitHub at https://github.com/shafaz89/EzrA_cell_division.

### Peptidoglycan extraction

*S. aureus* peptidoglycan was isolated as described (84). The concentration of extracted peptidoglycan was normalized based on colony-forming units (CFU), and samples were digested with mutanolysin (500 U/mL) for 18 h at 37°C. Following digestion, samples were neutralized with sodium hydroxide to pH 7.0, dried, and chemically reduced by the addition of 250 mM sodium borate and 3–5 mg of sodium borohydride. Reduction reactions were allowed to proceed for 30 min and were quenched by the addition of 20% phosphoric acid to pH 4.0, as described previously (84). Reduced muropeptides were separated by reversed-phase high-performance liquid chromatography (HPLC) using a C18 column (250 × 4.6 mm; ODS Hypersil, Thermo Scientific).

### Scanning electron microscopy (SEM)

Exponentially growing cells were fixed with 2.5% glutaraldehyde for 30 minutes and then washed three times with phosphate buffer (pH 7.2). The samples were dehydrated through a graded ethanol series (50–100%), critical-point dried with CO_2_, and sputter-coated with gold– palladium. SEM was performed at the Boston Electron Microscopy Center using a Hitachi S-4800 high-resolution field-emission scanning electron microscope (FE-SEM).

### LTA extraction and immunoblotting

LTA was extracted as described (85). Briefly, culture volumes were normalized based on CFU and lysed by bead beating. The lysates were centrifuged at 200 × g to remove glass beads, and the supernatants were further centrifuged at 16,000 × g for 15 min to pellet membranes and LTA. The resulting pellets were resolved by SDS-PAGE using 15% polyacrylamide gels and transferred to nitrocellulose membranes. Membranes were probed with an LTA-specific monoclonal antibody (mAb 55; Novus Biologicals) at a 1:200 dilution, followed by incubation with an HRP-conjugated anti-mouse IgG secondary antibody (Cell Signaling Technology). Immunoreactive bands were detected using SuperSignal chemiluminescent substrate (Thermo Scientific).

### WTA extraction and PAGE analysis

WTA was extracted as described (86). Briefly, CFU-normalized cells were washed once with 30 mL of buffer 1 (50 mM 2-[N-morpholino]ethanesulfonic acid [MES], pH 6.5) and resuspended in 30 mL of buffer 2 (4% [wt/vol] sodium dodecyl sulfate [SDS], 50 mM MES, pH 6.5). Samples were boiled and centrifuged at 10,000 × g for 10 min. The resulting pellet was sequentially washed with buffer 2, buffer 3 (2% NaCl, 50 mM MES, pH 6.5), and finally buffer 1. Pellets were treated with proteinase K, followed by washing with buffer 3 and multiple washes with distilled water. Extracted WTA was hydrolyzed with 0.1 M NaOH, and insoluble material was removed by centrifugation at 14,000 × g for 10 min. Supernatants were neutralized with 0.1 M acetic acid and extensively dialyzed against distilled H_₂_O prior to electrophoretic analysis. Polyacrylamide gels were prepared as previously described (86, 87). Electrophoresis was performed at 60 V for 24 h, at 4°C in Tris-Tricine running buffer (0.1 M Tris base, 0.1 M Tricine, pH 8.2) (88). WTA was visualized by sequential staining with alcian blue and silver (87).

### TUNEL assay

DNA breaks were detected using a TUNEL assay as described (89). Cells were fixed in formalin-free tissue fixative (Sigma-Aldrich), washed with PBS, and resuspended in GTE buffer (50 mM glucose, 20 mM Tris-HCl pH 7.5, 10 mM EDTA). To permeabilize the cell wall, cells were briefly treated with lysostaphin (2.5 µg/mL). DNA breaks were then labeled using the In Situ Cell Death Detection Kit, Fluorescein (Roche) according to the manufacturer’s instructions. Cells were washed with PBS and imaged by fluorescence microscopy.

**Table 1:**
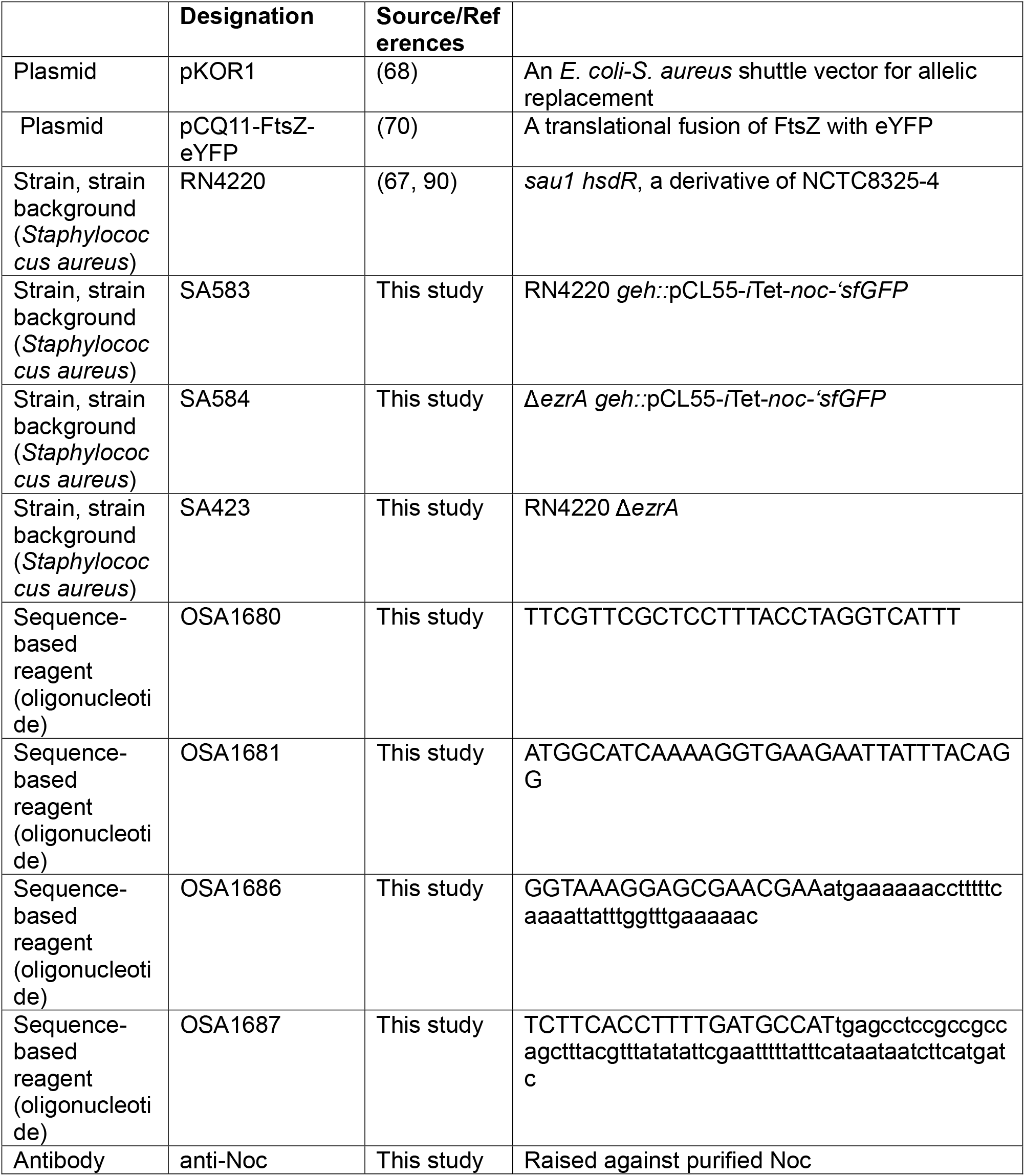
Key resources table.

## Acknowledgements

We thank members of the laboratory for helpful discussions and advice throughout this project. We are especially grateful to Derek Elli for his assistance. We thank Drs. Dora Bonini, Mariana Tinajero-Trejo, and Simon Foster (University of Sheffield) for generously sharing plasmids, and Dr. Shirin Kaboli and William Fowle (Boston Electron Microscopy Center) for their help acquiring the SEM images. This work was supported by grant AI038897 from the National Institute of Allergy and Infectious Diseases.

